# A Proposed Statistical Approach for Conducting a Longitudinal Assessment of Circulating Tumor DNA

**DOI:** 10.1101/2025.04.03.646973

**Authors:** Christopher R. Pretz, Jiemin Liao, Caroline Weipert, Leslie Bucheit, Leylah Drusbosky, Amar K. Das

## Abstract

As circulating tumor DNA (ctDNA) may reflect cancer progression, understanding its temporal evolution can inform clinical decision-making in precision oncology. Temporal changes in ctDNA often exhibit complex patterns, varying significantly between and within patients. Factors such as patient characteristics and treatment regimens can further impact these changes. Traditional statistical methods may fall short in adequately characterizing patient-level ctDNA evolution over time, highlighting the need for advanced approaches. In this study we provide a framework to guide the identification of optimal models for the analysis of genetic biomarkers in liquid biopsy settings. Specifically, we focus on hierarchical mixed-effects models as they provide both cohort and patient-level insights. To illustrate the versatility of these models, we conduct an exploratory analysis of a real-word data consisting of patients with advanced colorectal cancer. In our analysis, we discovered that a hierarchical linear spline mixed-effects model was most optimal. Based on this finding, we used the model to generate cohort and patient-level response patterns, where patient-level results compare ctDNA trajectories using different combinations of patient demographics, medical history, treatments regimen, and outcomes. Finally, we discuss how results could potentially assist in constructing a patient monitoring system to help inform patient care.

## INTRODUCTION

Plasma-based next generation sequencing for comprehensive genomic profiling of oncology patients with advanced solid tumors is recommended by numerous guidelines and expert consensus statements including those from the National Comprehensive Cancer Network, the American Society of Clinical Oncology, and the European Society for Medical Oncology. Plasma-based ‘liquid biopsies’ have many clinical applications in advanced cancers including genotyping to identify of actionable biomarkers with targeted therapeutic options and/or resistance mechanisms, monitoring of therapy response, and prognostication. Liquid biopsies have also been applied in cancer screening and early detection, detection of molecular residual disease after curative intent treatment, and predicting patients at risk of recurrence.^1,2^ Additional studies suggest serial measurements of circulating tumor DNA (ctDNA) may provide insight into disease progression. For instance, Sanz-Garcia et al. discussed how temporal ctDNA can have widespread utility in disease management across cancer types, while McLaren et al. proposed serial ctDNA testing be used to inform radiotherapy dosing.^3,4^ In addition to illustrating how serial ctDNA testing can support disease management, these studies highlight the promise of providing a custom-tailored patient-level approach to oncology.

Interpreting serial ctDNA results can be challenging. ctDNA levels often vary substantially within and between patients, posing distinct obstacles when analyzing these types of data.^4^ Similarly, because patient characteristics (such as age, disease characteristics, etc.) can impact temporal ctDNA patterns, these factors should be accounted for in analyses.^5,6^ To analyze complex temporal ctDNA patterns while simultaneously considering how patient characteristics associate with these patterns, we propose the adoption of a family of hierarchical linear, non-linear, and spline mixed-effects models (HMEM). The foundations of these models are well established and have been widely discussed and applied.^7–17^ As these prior works point out, this family of models has many advantages in comparison to traditional longitudinal analyses such as use of change scores, calculating ratios between baseline measures and future time points, or response profile analysis. The main expected benefits of using HMEMs to analyze serial ctDNA are: (1) the ability to capture cohort and patient-level biomarker evolution, (2) patient characteristics, treatment information, and patient outcomes are readily incorporated, (3) time points between measures need not be equally spaced and missing data is less problematic, and (4) cohort and patient-level longitudinal projections can be generated.

To our knowledge, the implementation of HMEMs in the analysis of complex longitudinal genetic data has been previously unaddressed. As such, this study aims to introduce and provide examples of how HMEMs can be used to achieve a comprehensive understanding of continuous or pseudo continuous longitudinal genetic biomarkers (although generalized versions of HMEMs that support the analysis of binary, ordinal etc. response variables exist ^7-11^). We begin by familiarizing the reader with a basic HMEM followed by a discussion of more complicated models. Next, an approach for identifying the most optimal HMEM is presented based on the analysis of a retrospective real-world cohort of patients diagnosed with advanced colorectal cancer (CRC). Once an optimal model is identified, we provide several examples of how the proposed analytic approach can be used to conduct an extensive exploratory analysis and conclude with a discussion how model results can be applied to patient monitoring.

## METHODS

### Patient Cohort and Data Source

In this study we illustrate the application of a HMEM by analyzing a retrospective cohort of patients diagnosed with advanced CRC who received chemotherapy and/or targeted therapies. This cohort is based on real-world observational data obtained from the GuardantINFORM database, which includes anonymized clinical-genomic data and structured commercial payer claims from both inpatient and outpatient facilities across academic and community settings. Selected patients had at least three Guardant360 (G360) liquid biopsy tests ordered by US healthcare providers between June 2014 and June 2024. The generation of de-identified data sets by Guardant Health for research purposes was approved by the Advarra Institutional Review Board; patient identity protection was maintained throughout the study in a de-identified database. Retrospective analysis of de-identified data is approved by Advarra IRB number *Pro00034566*. All patients were required to have at least one blood sample within 90 days prior to initiation of chemotherapy or targeted therapies, and at least two blood samples while on treatment, or within 30 days post end of line of therapy. For patients with multiple lines of therapy meeting these criteria, the earliest line of therapy was selected for study inclusion. Finally, patients with suspected incidental germline mutations were removed from the cohort. The date of first G360 test was defined as the index date. The GuardantINFORM database is fully de-identified and complies with Sections 164.514 (a)–(b)1ii of the US Health Insurance Portability and Accountability Act (HIPAA) regarding the determination and documentation of statistically de-identified data. Given that the data is de-identified, obtaining patient consent was waived by an external Institutional Review Board.

### Response Variable and Study Covariates

Although the proposed modeling approach can be used to analyze other complex longitudinal biomarkers, our response variable, ctDNA levels, is defined as the maximum proportion of somatic variant allele frequency in plasma cell-free DNA, as detected through liquid biopsy. In cases where ctDNA levels fall below the assay’s limit of detection, values were replaced with ctDNA levels of 0.02%, consistent with the lowest value in the cohort at the time of analysis. To assess the impact of this imputation, a sensitivity analysis was performed using replacement values ranging from 0.007% to 0.07%, which found that replacement values had a minimal impact on overall results. All covariates except mortality (alive vs deceased) were captured at baseline, defined as 183 days prior to the index date. Baseline covariates include age (in years), smoking status (yes/no), gender (female/male), treatment type (chemotherapy/targeted) and the Charlson Comorbidity Index (CCI). Data were extracted using SAS software package 9.4, and all statistical analysis was performed using R version 4.1.3.

### Statistical Approach

In this section, we introduce select HMEMs, highlight their utility, and explain the process used to identify the model that best fits the response variable. Our comparator pool includes various hierarchical linear, non-linear, and spline mixed-effects models. While it is not possible to provide an in-depth discussion of each model, a summary of the HMEMs considered is provided in Table 1.0. We begin by presenting the mathematical details of a linear HMEM and then build upon this exposition to discuss a cubic spline HMEM. Due to the hierarchical structure of the models, each is comprised of two levels of equations. The level-1, or patient-level equations, capture the temporal patterns for each patient. For linear models, we consider those that contain linear, quadratic, and cubic terms. Consequently, the level-1 equation for a linear HMEM is expressed as follows:

**Table 1.0.**
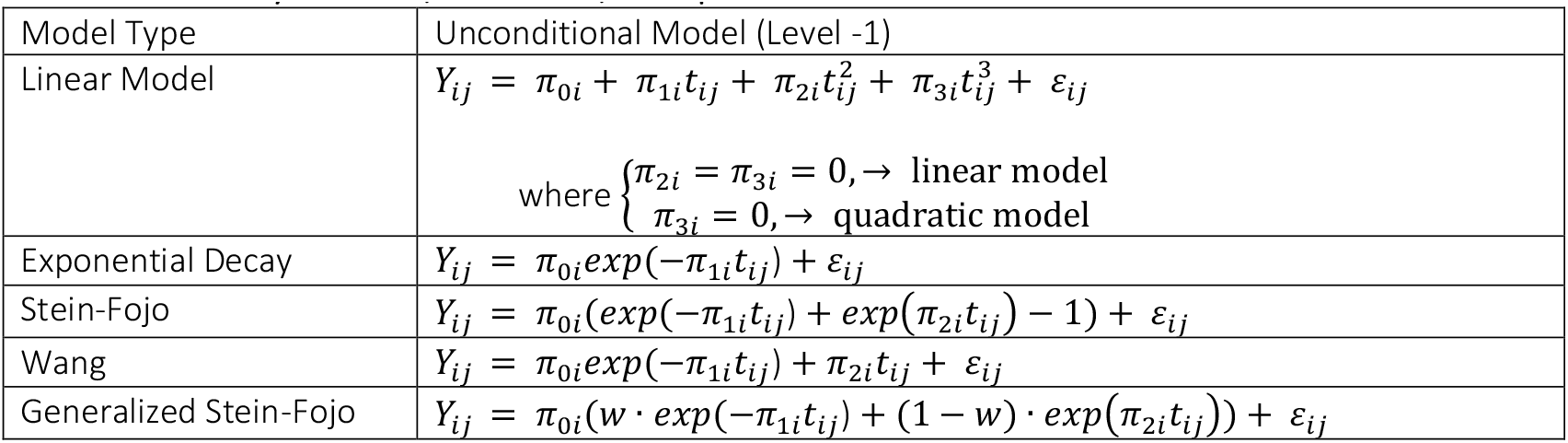
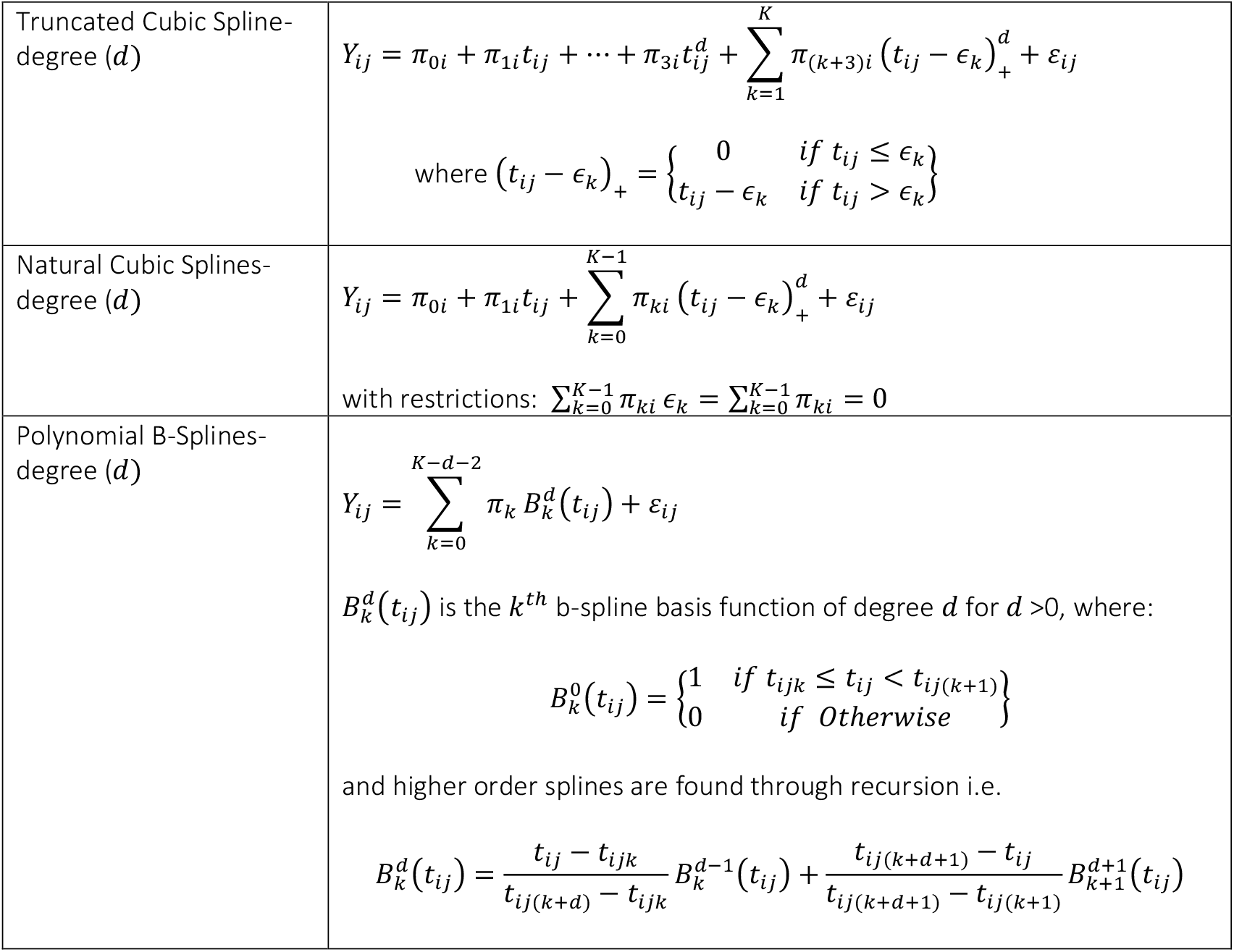
Summary of Linear, Non-linear, and Spline Level-1 Models.

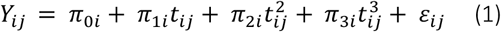

In this equation, the ctDNA measurements (or a transformation thereof) captured over time are represented by the *Y*_*ij*′*S*_, where *i* indexes the patient and *j* indexes the measurement occasion. Timepoints captured within the patient are given by *t*_*ij*_ and the 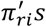 represent the *r* = 4 growth parameters. Here, π_0*i*_ denotes the intercept, π_1*i*_ represents the linear term, and π_2*i*_ represent the quadratic term etc. The last term in equation (1), ε_*ij*_, represents the model error and is assumed to be normally distributed with a mean of zero and a variance of σ^2^.

Next, we present the level-2, or cohort-level equation, which links the growth parameters to study covariates, treatment types, and patient outcomes:

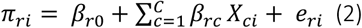

where, *X*_*ci*_ represents the covariate and *C* denotes the number of covariates ranging from 1 to *C*. As is typical in hierarchical models, numerical covariates are centered about their respective means. ^18^ Additionally, *β*_*r*0_ is the intercept for each corresponding π_*ri*_, and *β*_*rc*_ captures the linear relationship between a given growth parameter and covariate, where a unique set of *β*_*rc′S*_ can be specified if desired. When a model is free of covariates (i.e. π_*ri*_ = *β*_*r*0_) it is referred to as an unconditional model—otherwise—it is referred to as a conditional model. Finally, *e*_*ri*_ represents a random component and is assumed to follow a multivariate normal distribution:

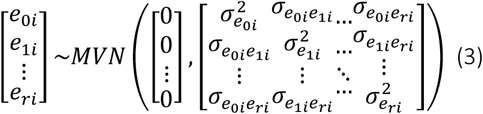

A key strength of the HMEM is its ability to generate results based upon the desired output. These results assume the form of a trajectory of estimated temporal ctDNA values referred to as a response pattern. By inputting various combinations of covariate values, a wide array of patient-level response patterns is readily obtained. Similarly, if a cohort-level response pattern is desired, continuous covariates can be set to their cohort averages, and binary covariates can be set to their cohort proportions.

At this juncture, we shift our focus to a more complex HMEM, one based on a truncated cubic spline, which is a natural extension of a linear model.^19^ A mathematical representation of a level-1 truncated cubic spline is given by:

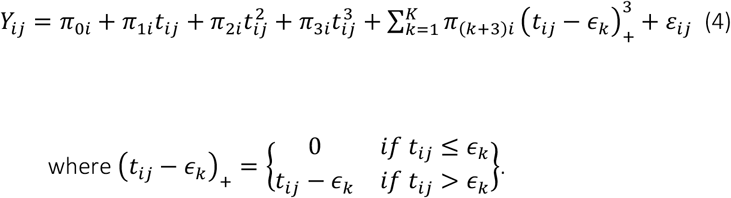

The primary distinction between equations (1) and (4) lies in the incorporation of the summation term. The summation term contains the *K* knot values given by *ϵ*_*k*_, where *ϵ*_1< … <_ *ϵ*_*k*_, along with an additional set of growth parameters i.e. π_(1+3)*i*,_ π_(2+3)*i*,…,_π_(*K*+3)*i*_. These growth parameters capture the relationship between the respective cubic polynomial terms (represented by 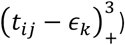 and the corresponding *Y*_*ij*_ values. Similar to the linear model, the level-2 equation for a truncated cubic spline is:

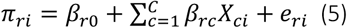

However, instead of the maximum of *r* = 4 growth parameters, equation (5) contains *r* = 4 + *K* growth parameters. While we specifically showcase a truncated cubic spline, other spline models such as a basis spline (often referred to as a B-spline) or natural cubic splines can also be used.^20–22^ Additionally, the degree (*d*) of a spline model can be modified. For instance, a level-1 truncated linear spline model (where *d* = 1) is given by:

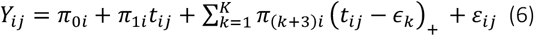

*A*long with various spline HMEMs, we also consider several non-linear models, including exponential decay, Stein-Fojo, Wang, and Generalized Stein-Fojo models.^23–29^ These non-linear models are typically used to describe changes in tumor volume over time—most often post treatment—but they can also capture ctDNA dynamics.^23^ Each non-linear model contains an exponential decay component, where the Stein-Fojo and Generalized Stein-Fojo models incorporate both exponential decay and exponential growth components.^26^ A summary of the models in our comparator pool is provided in Table 1.0.

An important property of the models summarized in Table 1.0 is they are differentiable, meaning the biomarker’s instantaneous rate of change (IRC) can be calculated. The IRC is primarily used for determining how the direction and speed of ctDNA levels change at a given point in time. As an example, the IRC for equation (4) can be determined by taking the first derivative with respect to time, resulting in:

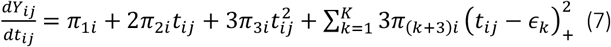

In this formula the IRC is given by the slope of the line tangent to the response pattern. Positive values correspond to an increasing IRC, negative values to a decreasing IRC, and IRC values of zero indicate either a peak or trough was reached, or that the trajectory is flat. The further the IRC value is from zero, the more extreme the rate of change is. As will be shown, the IRC can be used to create velocity plots, which graph the IRC as a function of time. Finally, the reduction in the residual variance gained from adding covariates can be calculated by extracting the variance estimates from the conditional and unconditional models and by evaluating the equation below.

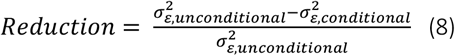

## STATISTICAL ANALYSIS and RESULTS

The study included 230 patients from the GuardantINFORM database who met the inclusion criteria. The average patient age was 58 years with 51% of the cohort being male. Additionally, 22% received targeted treatments, 21% were smokers, 35% became deceased during the follow-up period, and the average CCI is 1.1. The median study duration time was 613 days for surviving patients and 571 days for deceased patients. To satisfy model assumptions, ctDNA levels, originally expressed as a percent, were transformed into logits which mimic a continuous scale. Figure 1.0 presents histograms that show the logit transformation effectively mitigated the extreme skewness of the raw data. Likewise, the logit-adjusted spaghetti plot accentuates the variability in ctDNA levels over time, both within and between patients.

**Figure 1.0.**
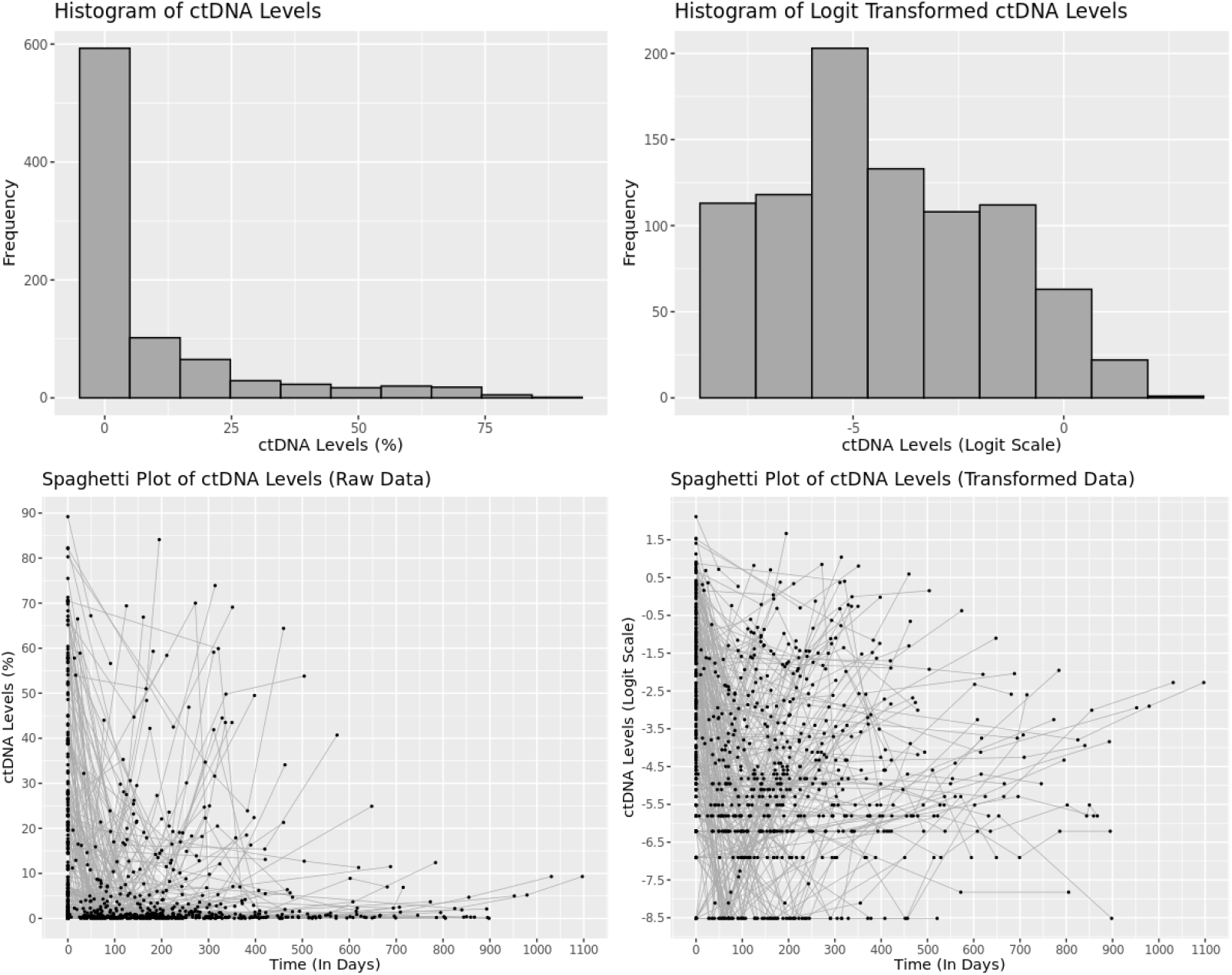
Distribution of ctDNA levels and logit transformed ctDNA levels along with corresponding spaghetti plots.

Guided by the equations in Table 1.0, we analyzed the CRC cohort using the series of models listed in Table 2.0. Our initial focus was on level-1 models, as we aimed to identify which model fit the patient-level data best. Model performance was evaluated using the Akaike information criteria (AIC) and Bayesian information criteria (BIC), with smaller values indicating more optimal models. Once an optimal model was identified, covariates were incorporated to enhance predictions and/or serve as statistical controls. Since spline models can be parametrized using various knot solutions, models containing one to four internal knots were investigated with knot placement determined by data quantiles. Furthermore, linear, quadratic, and cubic splines (i.e. *d* = 1, 2, *and* 3) were also considered. To keep the size of the model pool manageable, for truncated and natural splines, we limit our investigation to those containing third order polynomials. The model description, and the corresponding AIC and BIC values, are reported below.

**Table 2.0.** Unconditional models with corresponding information criterion values.

| Unconditional Model | AIC | BIC |
| --- | --- | --- |
| Linear Model ($\pi_{2i} = \pi_{3i} = 0$) | 3968.19 | 3996.822 |
| Linear Model ($\pi_{3i} = 0$) | 3925.369 | 3973.089 |
| Linear Model | 3869.706 | 3941.285 |
| Exponential Decay | 4115.82 | 4144.451 |
| Stein-Fojo | 3909.995 | 3957.714 |
| Wang | 4030.574 | 4078.293 |
| Generalized Stein-Fojo | DNC | DNC |
| B-spline ($d = 1$, knots = 1) | 3832.925 | 3880.644 |
| B-spline ($d = 1$, knots = 2) | 3694.092* | 3765.671* |
| B- spline ($d = 1$, knots = 3) | 3712.607 | 3812.818 |
| B-spline ($d = 1$, knots = 4) | 3699.717 | 3833.332 |
| B-spline ($d = 2$, knots = 1) | 3925.369 | 3973.089 |
| B-spline ($d = 2$, knots = 2) | 3743.848 | 3815.427 |
| B- spline ($d = 2$, knots = 3) | 3710.601 | 3810.811 |
| B-spline ($d = 2$, knots = 4) | 3708.217 | 3841.831 |
| B-spline ($d = 3$, knots = 1) | 3869.466 | 3941.046 |
| B-spline ($d = 3$, knots = 2) | 3869.466 | 3941.046 |
| B- spline ($d = 3$, knots = 3) | 3707.84 | 3808.051 |
| B-spline ($d = 3$, knots = 4) | 3713.517 | 3847.131 |
| Truncated Cubic Spline (knots=1) | 3726.674 | 3826.885 |
| Truncated Cubic Spline (knots=2) | 3730.881 | 3864.495 |
| Truncated Cubic Spline (knots=3) | 3746.139 | 3917.929 |
| Truncated Cubic Spline (knots=4) | DNC | DNC |
| Natural Cubic Spline (knots=1) | 3917.486 | 3965.205 |
| Natural Cubic Spline (knots=2) | 3758.484 | 3830.063 |
| Natural Cubic Spline (knots=3) | 3703.155 | 3803.366 |
| Natural Cubic Spline (knots=4) | 3709.201 | 3842.816 |

DNC indicates model did not converge
\* Indicates Lowest AIC and BIC value

After evaluating the AIC and BIC values, we conclude that a linear spline with two internal knots fits the data best (Table 2.0).^30^ Parameter estimates for this model are provided in the supplemental material. Note that even though this model had the lowest information criteria, if a different dataset was analyzed, the best fitting model could easily assume the form of a different model.

Figure 2.0 maps out the progression of ctDNA levels over time at the cohort-level. The response pattern in Figure 2.0 reveals that across patients, ctDNA levels drop substantially between baseline and 60 days, then rise rapidly until 200 days, after which levels continue to increase, but do so at a less extreme rate. This response pattern is consistent with expected treatment effects—reflecting an initial treatment response followed by treatment resistance. The 95% confidence bands about the response pattern are also displayed, which shows increased model uncertainty as the number of datapoints decrease over time. Due to the flexibility of the spline model, a major advantage is that it reveals details hidden within the data that simpler models may not detect. Despite this advantage, the model does not account for the contingency that patients with different characteristics, treatments, and outcomes may exhibit different response patterns. To explore these contingencies, various conditional models were fit to the data. The first of these examines the difference between the response patterns for alive vs deceased patients after controlling for baseline age, gender, CCI, treatment regimen, and smoking status.

**Figure 2.0.**
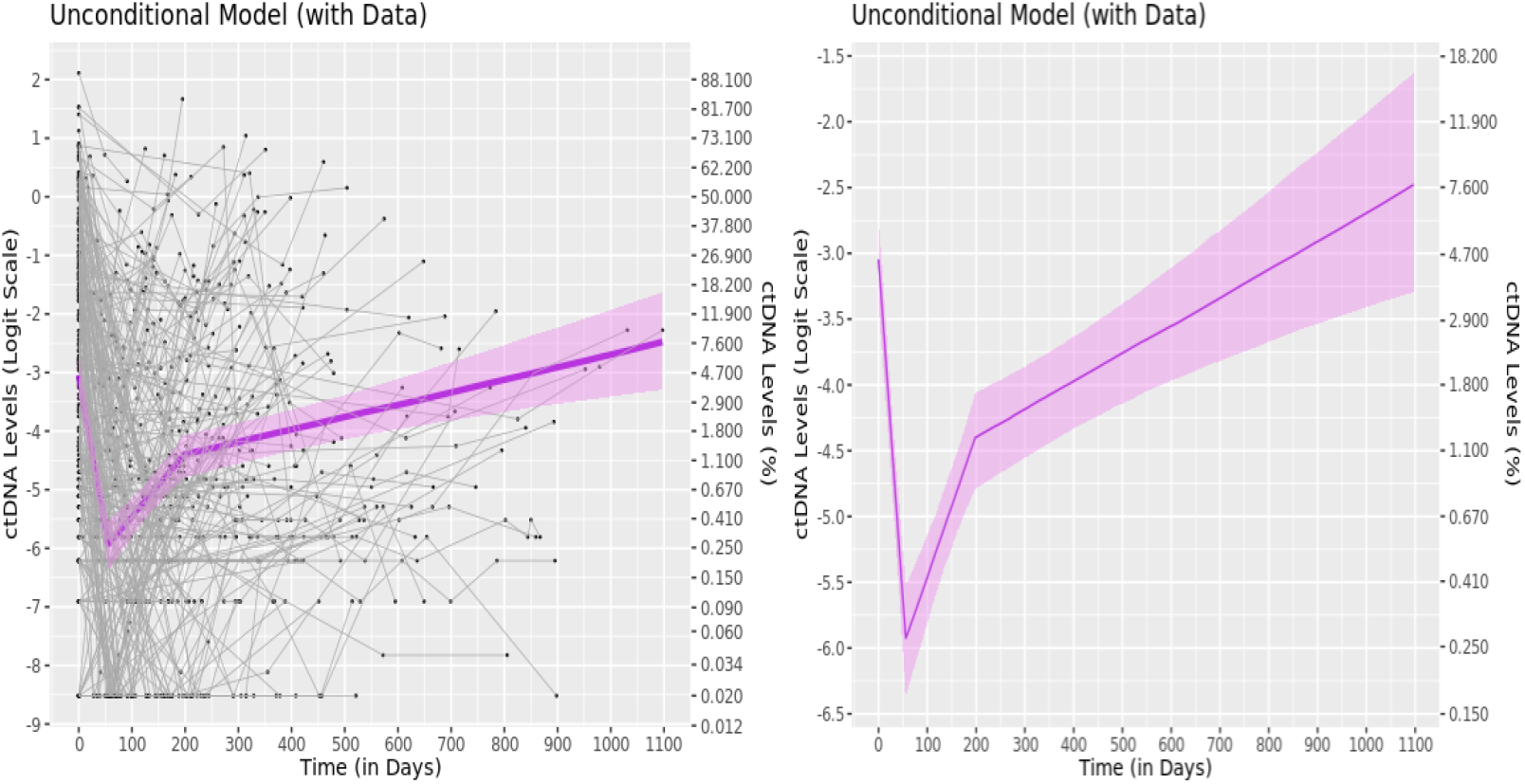
Unconditional model fit with and without datapoints for the CRC cohort. The violet function denotes the estimated response pattern for the cohort, while black dots connected by the gray lines indicates the change in ctDNA levels over time for each patient. The shaded violet region represents the 95% confidence bands for the estimated trajectory.

Figure 3.0 indicates that estimated baseline ctDNA values for patients who were alive (alive group) or became deceased (deceased group) during the follow-up period are nearly identical. Following baseline, estimates decrease for both groups, however, based on the velocity plot (see Figure 4.0), the decline is steeper in the alive group, with a drop of 0.0525 logits per day compared to 0.050 logits per day in the deceased group. By day 55 both groups reach their minimum ctDNA values: −6.1 logits (0.24%) for the alive group and −5.5 logits (0.41 %) for the deceased group. After day 55, estimated ctDNA levels rise for both groups, but the increase is more gradual in the alive group. The 95% confidence bands reveal that response patterns for the two groups begin to diverge around day 110. Shortly before day 200, both groups reach a local maximum: −4.9 logits (0.74 %) and −3.6 logits (2.7%) for the alive and deceased groups respectively. Beyond 200 days, estimated levels for each group increase at the same rate until at day 600 maximum ctDNA estimates are reached (Alive = −6.1 logits (0.24%), Deceased = −5.5 logits (0.41 %)). The analysis is limited to the first 600 days, as data beyond this point becomes sparce, leading to significant widening of the confidence bands.

**Figure 3.0.**
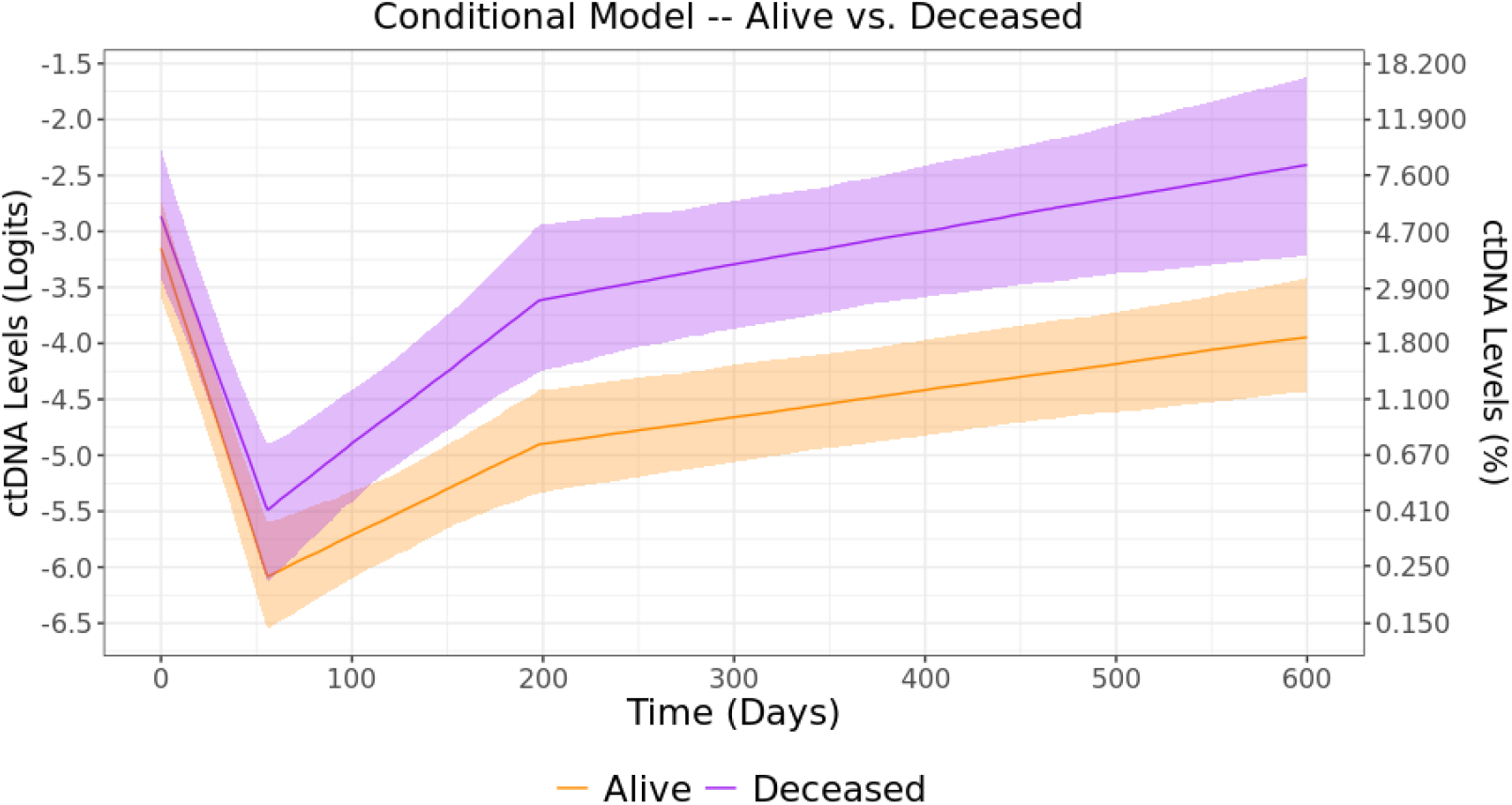
Conditional model—estimated response patterns with 95% confidence bands for Alive vs Deceased patients after controlling for baseline age, gender, CCI, treatment regimen, and smoking status.

**Figure 4.0.**
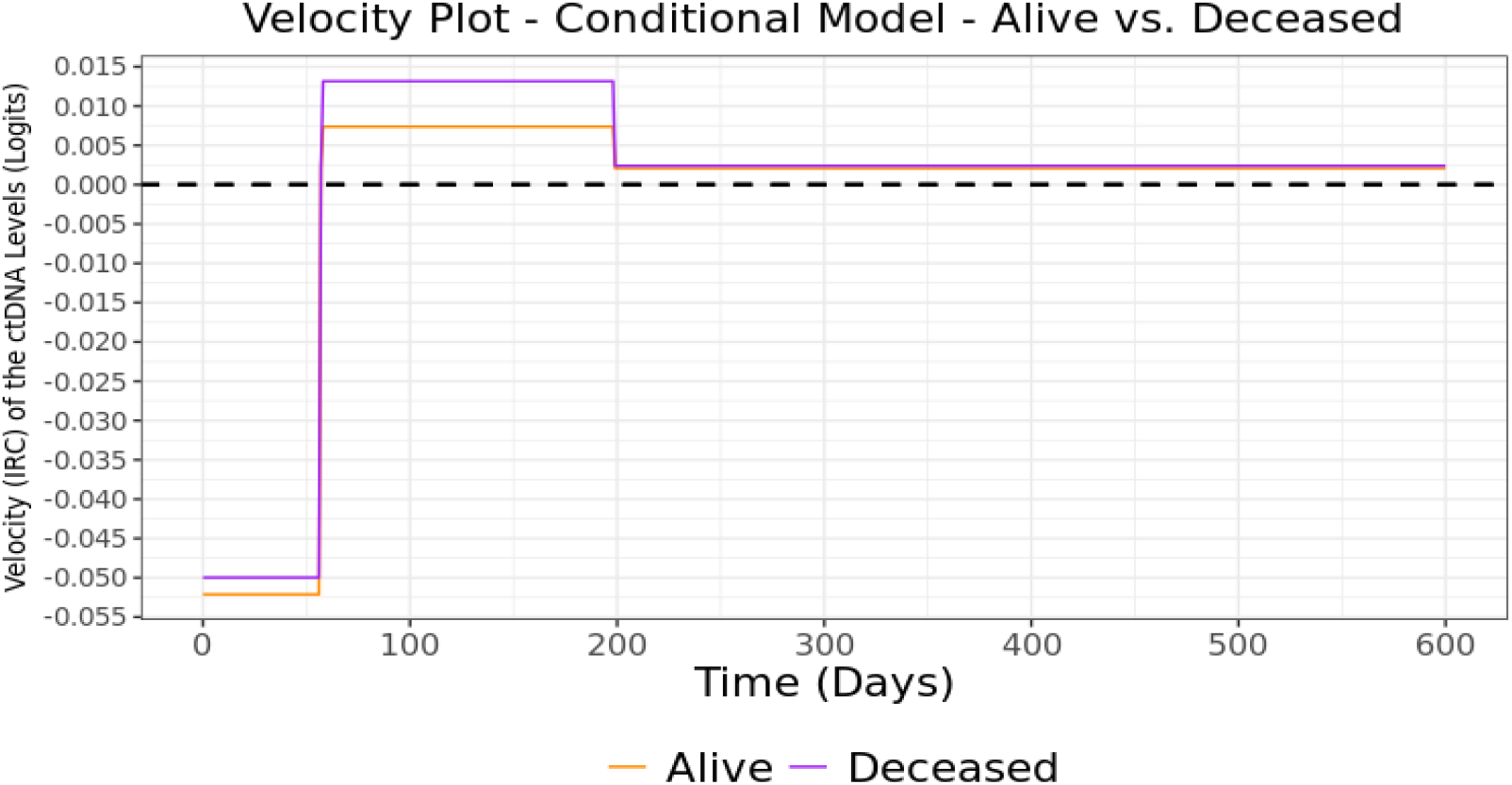
Velocity plot of conditional model—Alive vs Deceased patients after controlling for baseline age, gender, CCI, treatment regimen, and smoking status.

Another comparison of interest is whether response patterns differ between patients who received targeted therapies and those who underwent chemotherapy.

From inspecting Figure 5.0, the overlapping confidence bands suggest that, overall, no significant difference between the treatment regimens exist. Moving forward, we transition to exploring model projections between patients with different baseline characteristics, treatment regimens, and outcomes.

**Figure 5.0.**
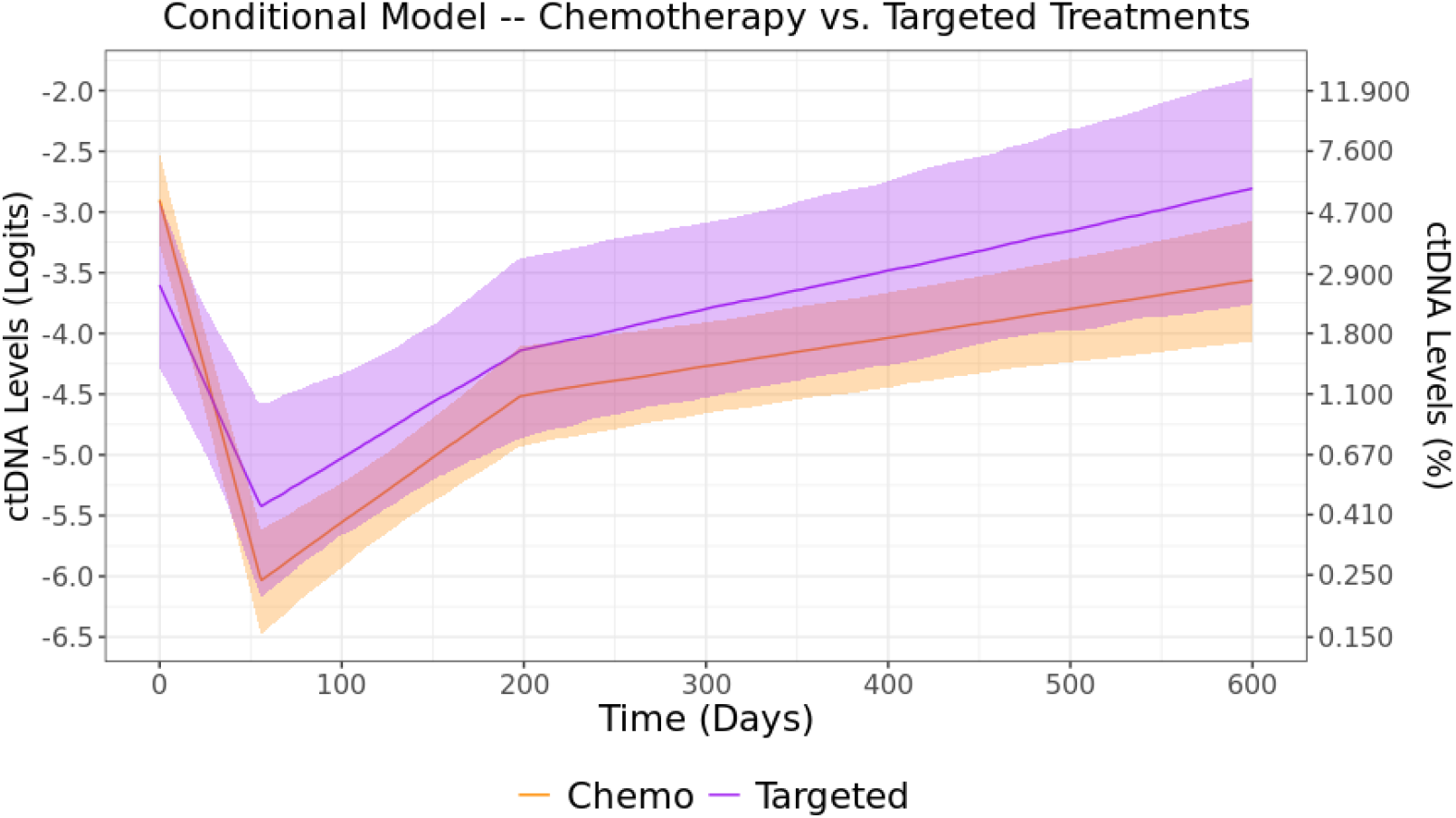
Conditional model—estimated response patterns with 95% confidence bands for Chemotherapy vs Targeted treatments after controlling for baseline age, gender, CCI, smoking status, and mortality.

Response patterns for each patient are illustrated in Figure 6.0, with the corresponding velocity plot shown in Figure 7.0. These response patterns depict the estimated average ctDNA levels over time for patients who share the same set of covariate values. For Patient 1, the covariate values are as follows: Baseline age 38-years, female, non-smoker, undergoing chemotherapy, with a CCI of 0, and alive throughout the study period. In contrast, Patient 2 is a 70-year-old male smoker who received a targeted treatment, had a CCI of 5, and was deceased during the study period. Model projections can be made for existing patients or can be extrapolated by inputting a reasonable combination of covariate values i.e. selecting continuous values that fall within their respective ranges, where in this example, response patterns are extrapolated. A comparison of these response patterns reveals that Patient 1 experiences a robust treatment response where Patient 2 does not. Also notable, is that Patient 1 has a higher estimated baseline ctDNA level, but by day 600, has a much lower value—a pattern expected for patients in the survivor group.

**Figure 6.0.**
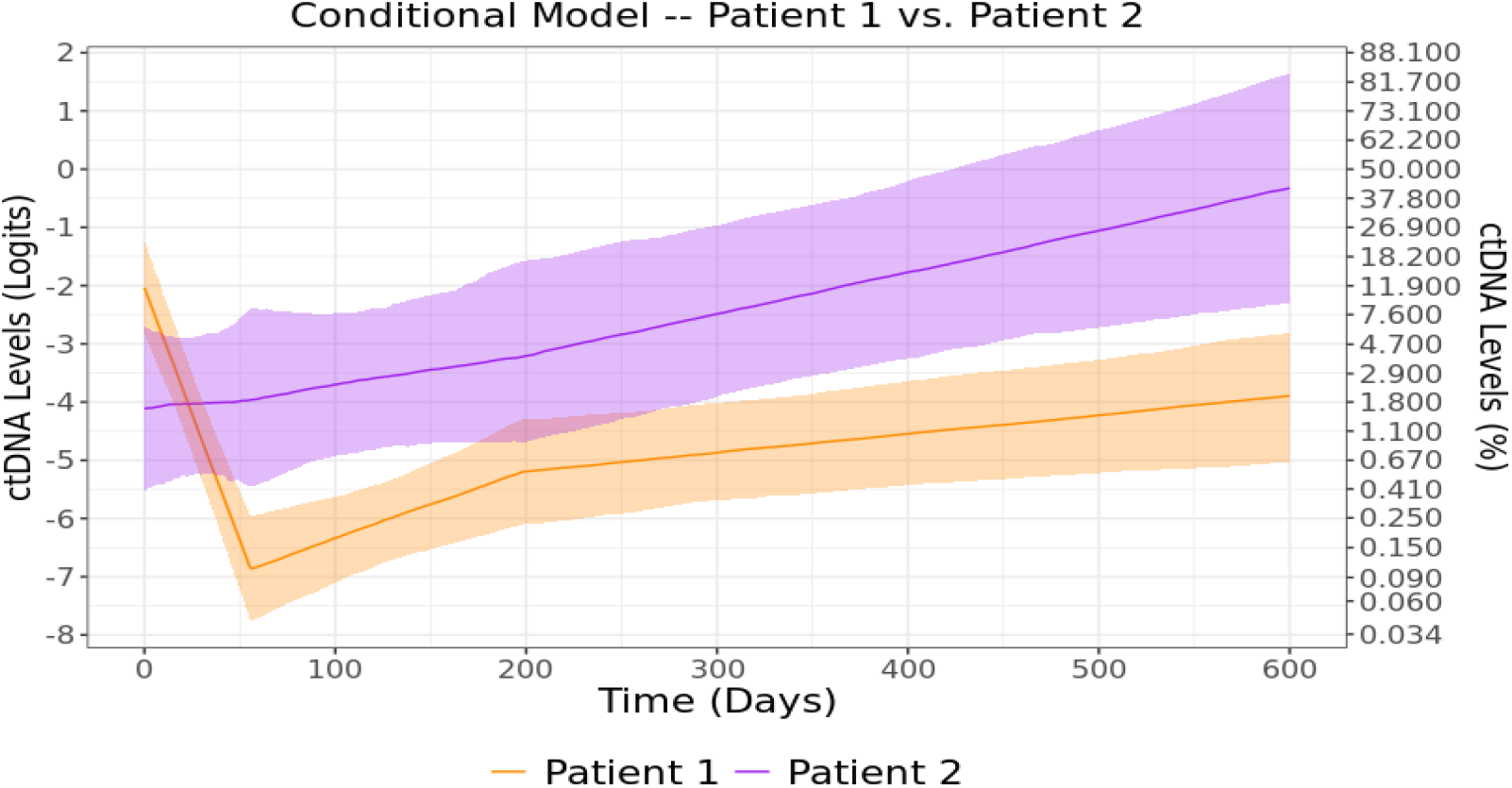
Conditional model— estimated response patterns with 95% confidence bands for Patient 1 vs Patient 2.

**Figure 7.0.**
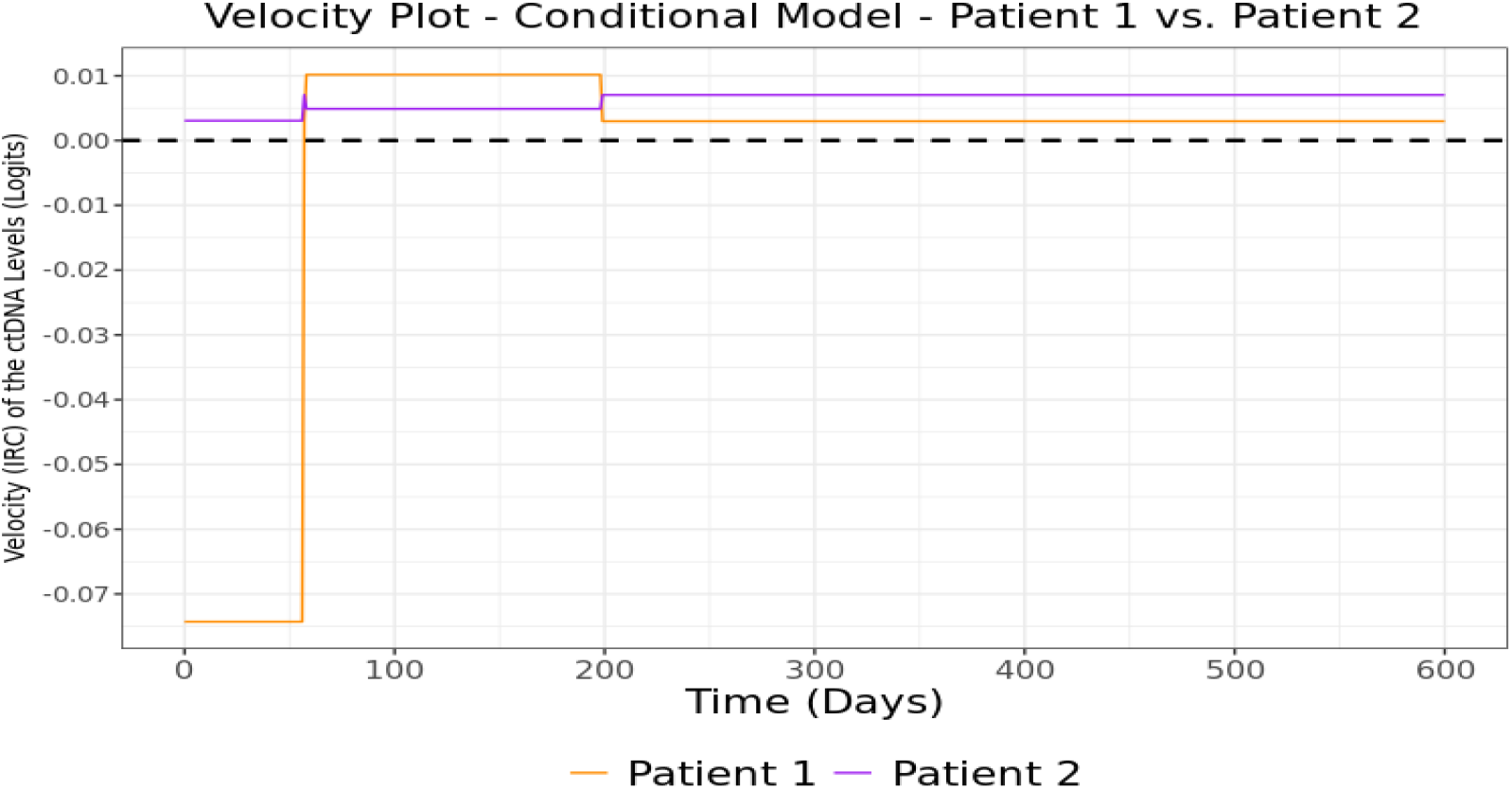
Velocity plot for conditional model—Patient 1 vs Patient 2.

We conclude our analysis by providing an example of how the model can be leveraged further to explore the data. For instance, the lack of a treatment response for Patient 2 raises questions. To investigate—a spectrum of scenarios—incrementally increasing in baseline age and CCI while holding the remaining covariates constant—are examined. These scenarios are summarized in Table 3.0, with the corresponding response patterns shown in Figure 8.0.

**Table 3.0.** Summary of Scenarios 1-4.

| Scenario 1 | Scenario 2 | Scenario 3 | Scenario 4 |
| --- | --- | --- | --- |
| Baseline Age = 60<br>Baseline CCI = 3 | Baseline Age = 70<br>Baseline CCI = 5 | Baseline Age = 75<br>Baseline CCI = 7 | Baseline Age = 80<br>Baseline CCI = 9 |

**Figure 8.0.**
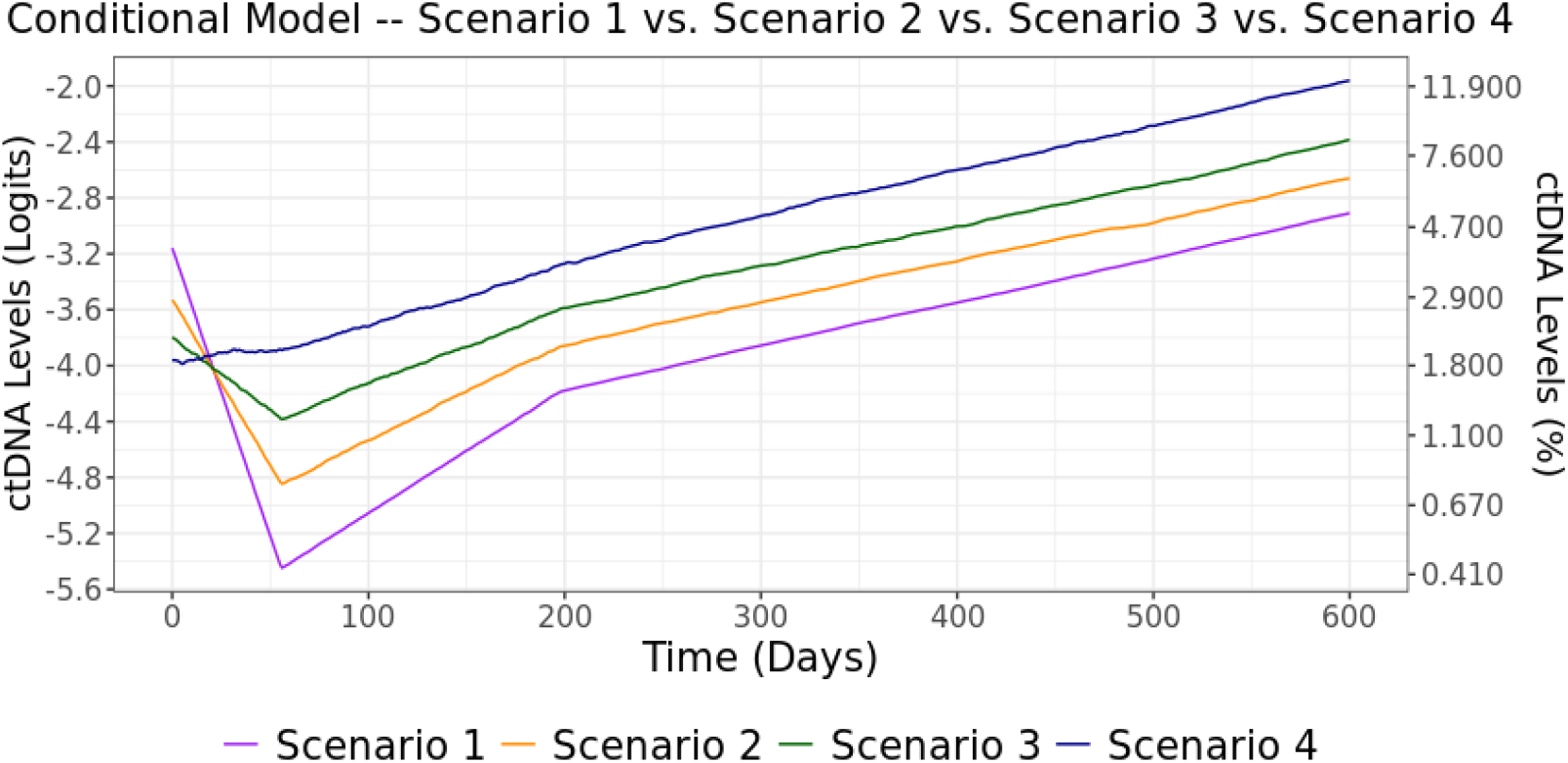
Estimated response patterns for scenarios 1-4.

The above response patterns suggest that, as baseline age and CCI increase, response to treatment appears to wane, implying that, in these data, for older patients and patients with greater comorbidity burden, treatment effect is attenuated.

## DISCUSSION

The purpose of this manuscript is to introduce a statistical approach to accommodate the analysis of complex longitudinal biomarker data where, to our knowledge, this approach has yet to be leveraged in a serial liquid biopsy genomic setting. As such, findings presented herein are not meant to represent bona fide results but instead are pedagogical in their intent. The proposed method is particularly useful when the desire is to obtain both cohort and patient-level results, for which HMEMs are well suited. As a result, linear, non-linear, and spline HMEMs are introduced and are recruited to construct a pool of candidate models. Of these candidates, a process to identify the model that fits the data best is outlined. To provide an example of optimal model identification, we analyzed a cohort of patients with advanced CRC based on real-world data. After ascertaining an optimal model, the model was used to conduct an exploratory analysis where versatility of our proposed approach is demonstrated by comparing various cohort and patient-level ctDNA response patterns. To complement these response patterns, velocity plots, which capture the biomarker’s IRC, are also generated. Since conducting an exploratory analysis is our intent, the 95% confidence bands serve as ‘guidelines’ in identifying substantial differences in ctDNA response patterns, where no attempt at statistical inference is made. Consequently, results presented are “data specific”, and therefore veracity of results should be established in other similar datasets, or, in future studies specifically designed to investigate impactful findings. The reader should be mindful that, in the examples provided, only a handful of results were investigated. This is because thousands of response patterns can be generated—each based on a unique combination of covariates. Thus, conducting a thorough investigation of the data would be daunting. Although we do not do so here, to alleviate this challenge, an RShiny, Excel, or a similar platform can be used to create an interactive tool that generates a response pattern for each covariate combination, allowing for the interrogation of large volumes of output.

Although we conducted an exploratory analysis based on observational data, this methodology is equally applicable to studies using representative cohorts or randomized control trials. When statistical inference is the goal controlling for type-I error and limiting the number of comparisons by focusing on predetermined hypotheses is essential. In these cases, hypotheses will likely be designed to compare response patterns between groups of interest but can also be based on conjecture regarding the nature of the relationship between response pattern behavior and the covariate values themselves.

Although we focus on using the statistical approach to achieve a compressive understanding of ctDNA dynamics, the proposed framework may also hold significant potential in precision oncology efforts, where assisting in patient monitoring is a potential clinical application. To illustrate, we provide a simple synopsis of how such a monitoring could be implemented. The core concept is that one can use an HMEM to construct patient-level response patterns, bifurcated by treatment response. In this instance these response patterns, and their corresponding confidence bands, can serve as a reference for patients with the same characteristics where the response pattern of a new patient can be compared against established patterns. If the pattern aligns (in the margin of error) with that of a responder, no intervention is necessary. However, if it mirrors a non-responder’s pattern action may be required. In sum, these model generated visualizations can help identify critical divergence points and offer insights into when intervention is warranted. Additionally, velocity plots, which can be used to compare IRC values, can enhance this process. However, before incorporating such a system into treatment decision making, rigorous internal and external model validation on the prediction of response variable is required. Furthermore, if survival based patient-level dynamic predictions are desired, the HMEM could be used as a longitudinal sub-model within a Joint modeling of longitudinal and time-to-even framework.^31^

We end by discussing some augmentations and limitations of the statistical approach. As response patterns are mathematical representations that leverage relationships between the covariates and the growth parameters, modifications of these relationships can be made to improve model performance. For instance, when the relationship between the response parameter and covariate(s) is non-linear, imposing a linear constraint may lead to an incorrect model specification. In such cases, a scatterplot can help to capture the true relationship and guide necessary adjustments. As with all models that capitalize on relationships between variables, such associations do not imply causality. Improving model efficiency, especially in the presence of multicollinearity, can often be achieved through model reduction, though the reduction process can be an arduous task due to the complicated nature of these models. Nevertheless, model reduction can be guided by utilizing the same information criteria discussed previously. Additionally, care should be taken when generating model projections. This is because an inadvertent use of nonsensical covariate combinations or covariate values outside the data range could produce nonsensical results. When timepoints are fully captured and equally spaced (not the case in our example), it becomes possible to account for the correlation structure of repeated measures. Correlation structures, such as first order autoregressive, compound symmetry, and spatial power structures can be explored, where the AIC or BIC can facilitate the identification of a suitable structure. Selecting an appropriate sample size is also an important consideration when conducting an analysis where minimum sample size recommendations for HMEMs are around 100 patients with at least 3 measurements per patient, though these models have been fit using sample sizes as low as 22 individuals.^32–34^ Finally, a number of ways in which k-fold cross-validation for mixed effects models has been proposed.^34^

## CONCLUSION

We present a framework that incorporates an assortment of HMEMs designed to capture a continuous prognostic genetic biomarker over time while incorporating covariate information. To demonstrate the potential of these models, we analyzed a real-world observational dataset of patients who have advanced CRC. This analysis provides a roadmap for identifying the most suitable model, where this model can be used to produce both cohort and patient-level results. With additional clinical validation and testing, the model may provide utility for patient monitoring and support clinical decision making in precision oncology practice.

## SOFTWARE

1. R Core Team (2022). R: A language and environment for statistical computing. R Foundation for Statistical Computing, Vienna, Austria. URL https://www.R-project.org/.
2. SAS Institute Inc., 100 SAS Campus Drive, Cary, NC 27513-2414, USA

## Acknowledgments

We thank Dr. Aaron Hardin, PhD for insights regarding ctDNA capture and the limit of detection of the G360 test.

## Author Contributions

C.R.P. conceived of the presented model and computational framework, analyzed the data, presented the results, and authored the manuscript with critical input from all co-authors. J.L. created the cohort, aided in the development of the model and computational framework, provided analytic support, and authored the manuscript. C.W., L.D., L.B., & A.D. provided critical content knowledge expertise, guidance, and input, and authored the manuscript.

## Data Availability Statement

The datasets generated during and/or analyzed during the current study are not publicly available and cannot be shared due to the use of a third-party healthcare claims database. Researchers interested in replicating our study or pursuing new research topics should contact Guardant Health (https://guardanthealth.com/products/biopharma-solutions/real-world-evidence/) directly.

## Additional Information

Competing Interests: All manuscript authors are employed by Guardant Health. Each receive an annually salary, bonus, and stock options that are commensurate with the author’s job description, experience, and level of education.

## SUPPLEMENTARY INFORMATION

**Supplemental Table 1.0.**
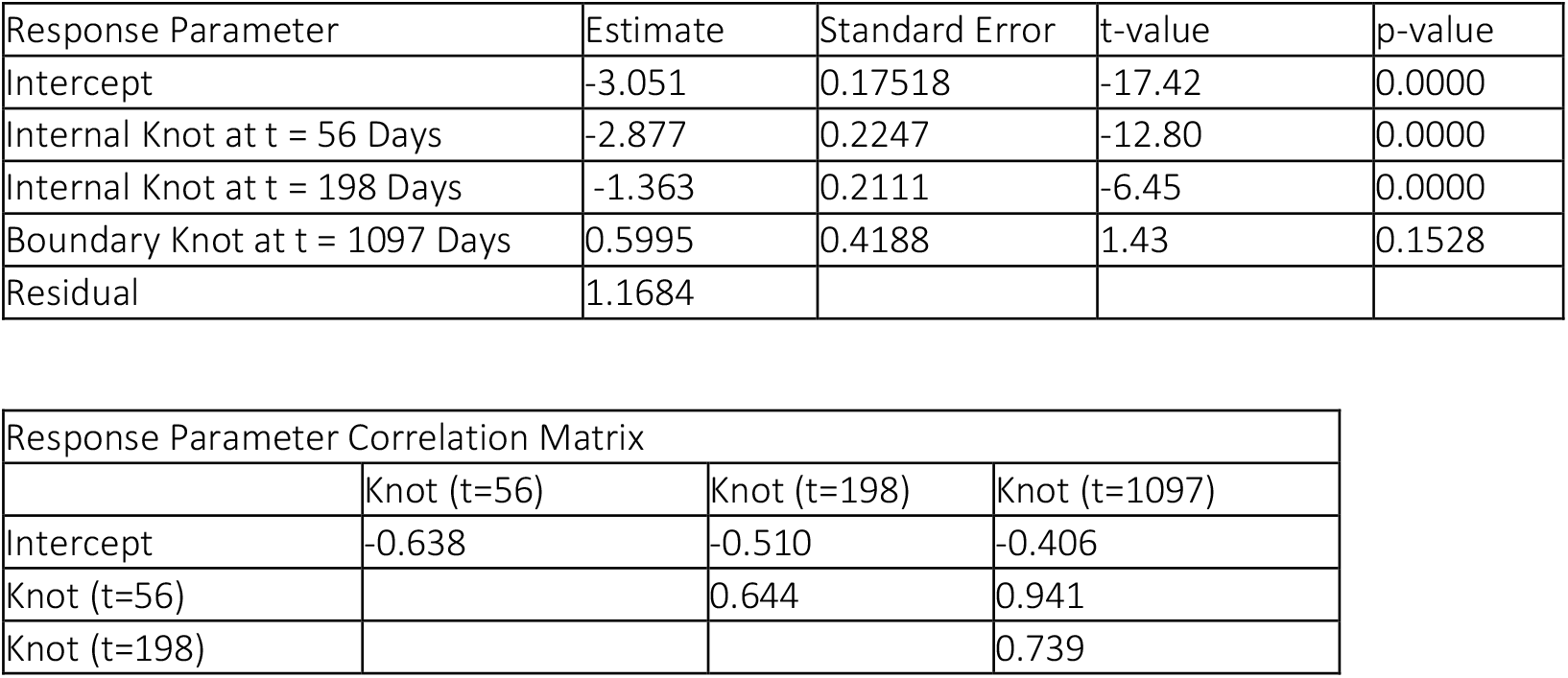
Summary of Best Fitting Unconditional Model.

| Response Parameter | Estimate | Standard Error | t-value | p-value |
| --- | --- | --- | --- | --- |
| Intercept | -3.051 | 0.17518 | -17.42 | 0.0000 |
| Internal Knot at t = 56 Days | -2.877 | 0.2247 | -12.80 | 0.0000 |
| Internal Knot at t = 198 Days | -1.363 | 0.2111 | -6.45 | 0.0000 |
| Boundary Knot at t = 1097 Days | 0.5995 | 0.4188 | 1.43 | 0.1528 |
| Residual | 1.1684 |  |  |  |

| Response Parameter Correlation Matrix |  |  |  |
| --- | --- | --- | --- |
|  | Knot (t=56) | Knot (t=198) | Knot (t=1097) |
| Intercept | -0.638 | -0.510 | -0.406 |
| Knot (t=56) |  | 0.644 | 0.941 |
| Knot (t=198) |  |  | 0.739 |

### Related Sample R Code

Below is sample R code for the HCSREM using a truncated cubic spline model (from the nlme package).^1^ In this sample code, we include 2 covariates and 3 knots. We present this code in a generic sense, where “data” could reference any longitudinal data expressed in the hierarchical fashion. The data for a truncated cubic spline must first be prepped, where the code to do so is provided below.

~~~
data$t = data$long_time_var
data$t2 = data$t^2
data$t3 = data$t^3
knot_1_time_point <- 50 #Time point of first knot
knot_2_time_point <- 150 #Time point of second knot
knot_1_time_point <- 250 # Time point of third knot
data$t3_knot_1=(data$t-knot_1_time_point)^3*I(data$t>knot_1_time_point)
data$t3_knot_2=(data$t-knot_2_time_point)^3*I(data$t>knot_2_time_point)
data$t3_knot_3=(data$t-knot_3_time_point)^3*I(data$t>knot_3_time_point)
~~~

### The R code for the HCSREM using a truncated cubic spline

~~~
HCSREM_TCS <- nlme(response_var ~ Int + B_lin*t + B_quad*t2 + B_cube*t3 + B_knot_1*t3_knot_1 + B_knot_2*t3_knot_2 + cov1_int*cov1 + B_lin_cov1*cov1*t + B_quad_cov1*cov1*t2 + B_cube_cov1*cov1*t3
+ B_knot_1_cov1*t3_knot_1*cov1 + B_knot_2_cov1*t3_knot_2*cov1 + B_knot_3_cov1*t3_knot_3*cov1 + cov2_int*cov2 + B_lin_cov2*cov2*t + B_quad_cov2*cov2*t2 + B_cube_cov2*cov2*t3 + B_knot_1_cov2*t3_knot_1*cov2 + B_knot_2_cov2*t3_knot_2*cov2 + B_knot_3_cov2*t3_knot_3*cov2,
                             data=data,
                         fixed= Int + B_lin + B_quad + B_cube + B_knot_1 + B_knot_2 + B_knot_3 +
cov1_int + B_lin_cov1 + B_quad_cov1 + B_cube_cov1 + B_knot_1_cov1 +
B_knot_2_cov1 + B_knot_3_cov1 + cov2_int + B_lin_cov2 + B_quad_cov2 + B_cube_cov2 +
B_knot_1_cov2 + B_knot_2_cov2 + B_knot_3_cov2 ~ 1,
                           random= Int + B_lin + B_quad + B_cube + B_knot_1 + B_knot_2 + B_knot_3 ~ 1,
groups= ~ Patient_id,
start= c(0, 0, 0, 0, 0, 0, 0, 0, 0, 0, 0, 0, 0, 0), method = ‘ML’)
~~~

### Below is sample R code for the HCSREM using a natural cubic spline model. In this sample code, we include 2 covariates and 3 knots

~~~
HCSREM_NCS <- lme(response_var ~ cov1*ns(long_time_var, df = 4) + cov2*ns(long_time_var, df = 4), random = ~ ns(long_time_var, df = 4) | Patient_id, data =data, method =‘ML’)
~~~

Below is sample R code for the HCSREM using a cubic B-spline model. In this sample code, we include 2 covariates and 3 knots.

~~~
HCSREM_NCS <- lme(response_var ~ cov1*bs(long_time_var, df = 4) + cov2*bs(long_time_var, df = 4), random = ~ bs(long_time_var, df = 4) | Patient_id, data =data, method =‘ML’)
~~~

*The all of the model statements in the above code can be expanded to accommodate additional knots and additional covariates as needed.

Note that using nlme() provides more coding flexibility in comparison to using lme(), but constructing the code can be much more complicated. As the dataset we used is not publicly available, this code can be applied many types of hierarchical longitudinal data. We suggest using the *prothro* dataset presented in the JMBayes2 R package as it is a good example of a suitable datasest.^2^

1. Pinheiro J, Bates D, R Core Team (2023). *nlme: Linear and Nonlinear Mixed Effects Models*. R package version 3.1-164, https://CRAN.R-project.org/package=nlme.
2. Rizopoulos D, Papageorgiou G, Miranda Afonso P (2023). _JMbayes2: Extended Joint Models for Longitudinal and Time-to-Event Data_. R package version 0.4-5, https://CRAN.R-project.org/package=JMbayes2.

